## Supplementary Figures for "Kaposi’s sarcoma-associated herpesvirus induces specialised ribosomes to efficiently translate viral lytic mRNAs"

DIMT1

a

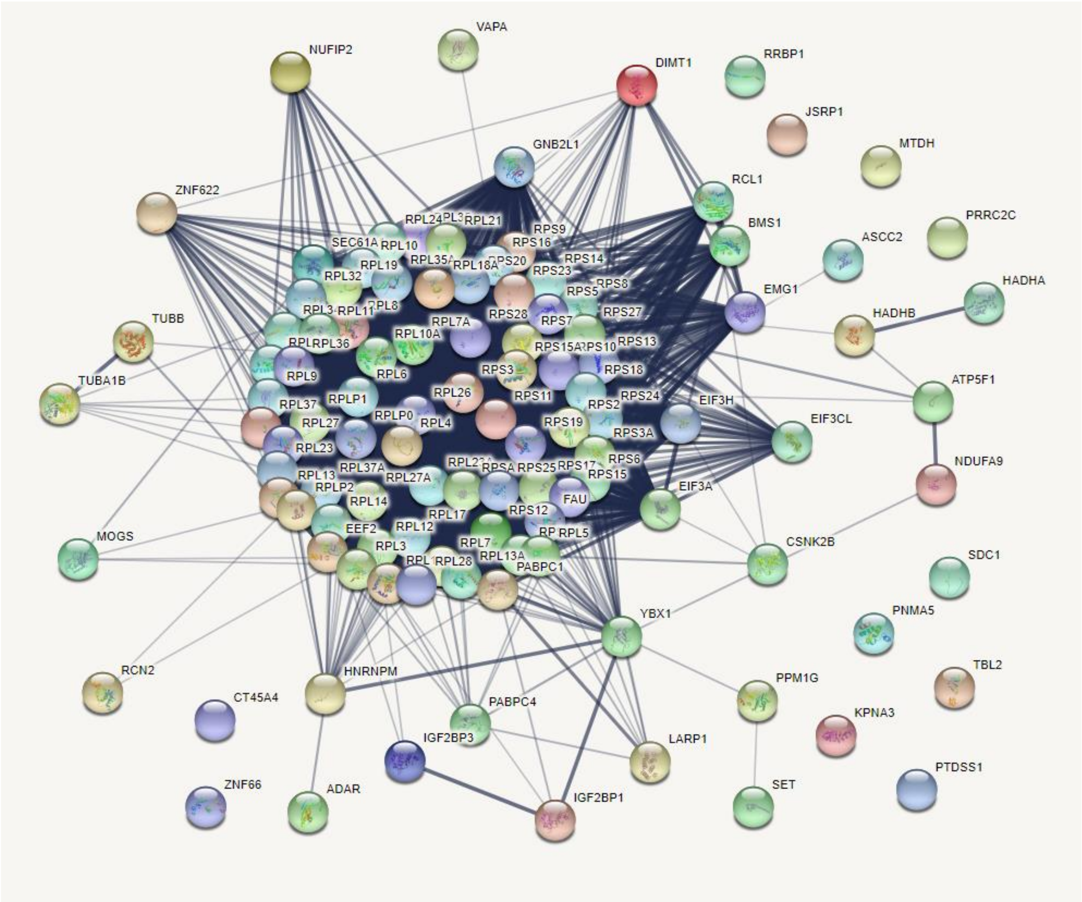


b

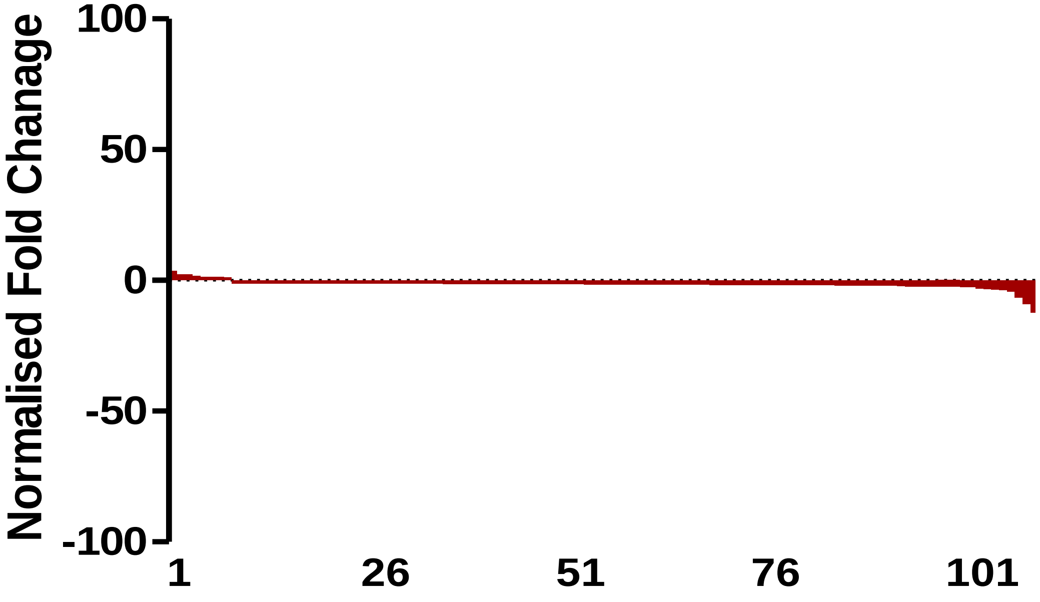


**Extended Data Figure 1. TMT LC-MS/MS analysis of DIMT1 Twin-Strep-tag® pulldowns from a latent and 24 hour post lytic reactivation TREx BCBL1-Rta cell line.** STRING protein interaction map of identified proteins **(a)**. Fold change of all interacting proteins from latent compared to 24 hours post lytic reactivation **(b)**.

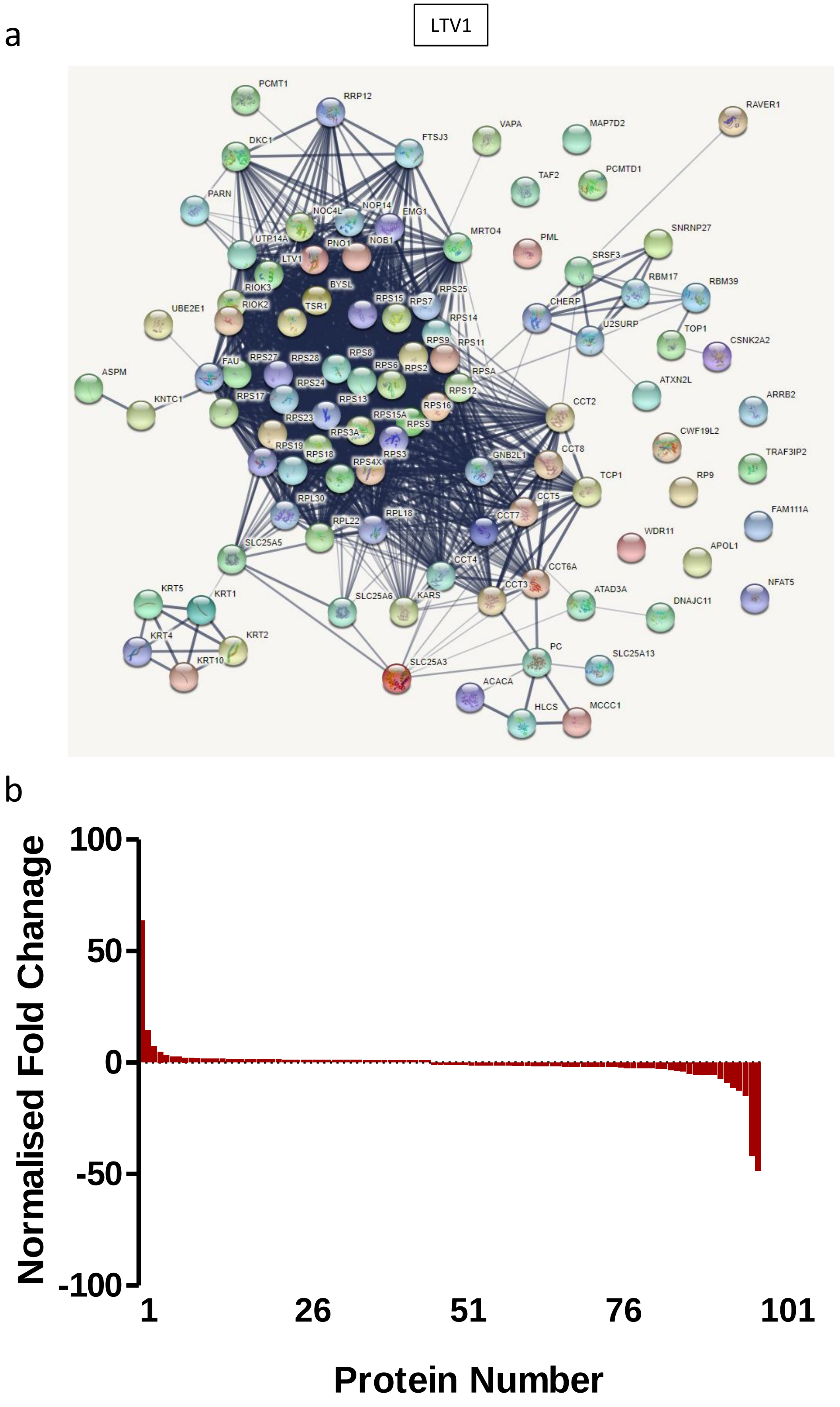


**Extended Data Figure 2. TMT LC-MS/MS analysis of LTV1 Twin-Strep-tag® pulldowns from a latent and 24 hour post lytic reactivation TREx BCBL1-Rta cell line.** STRING protein interaction map of identified proteins **(a)**. Fold change of all interacting proteins from latent compared to 24 hours post lytic reactivation **(b)**.

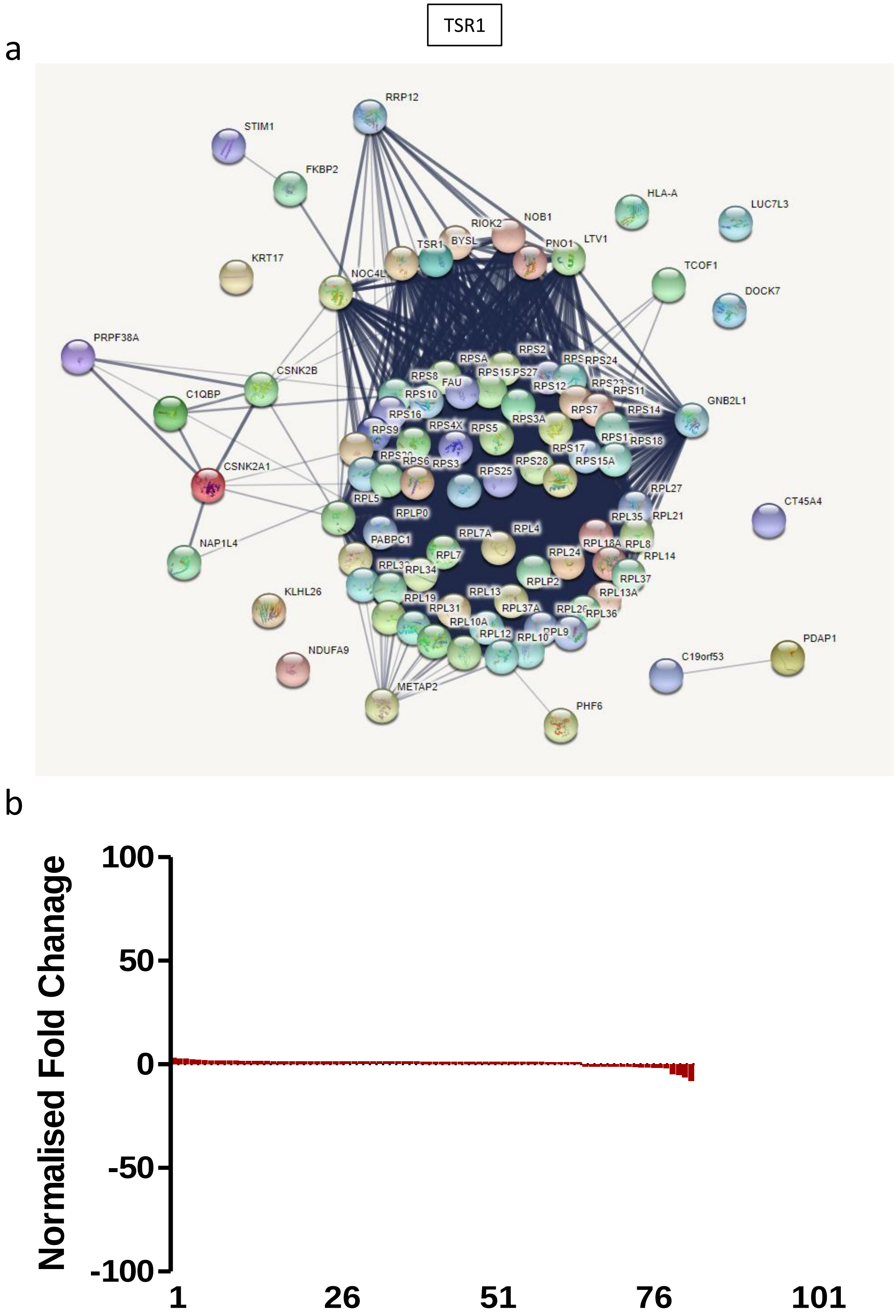


**Extended Data Figure 3. TMT LC-MS/MS analysis of TSR1 Twin-Strep-tag® pulldowns from a latent and 24 hour post lytic reactivation TREx BCBL1-Rta cell line.** STRING protein interaction map of identified proteins **(a)**. Fold change of all interacting proteins from latent compared to 24 hours post lytic reactivation **(b)**.

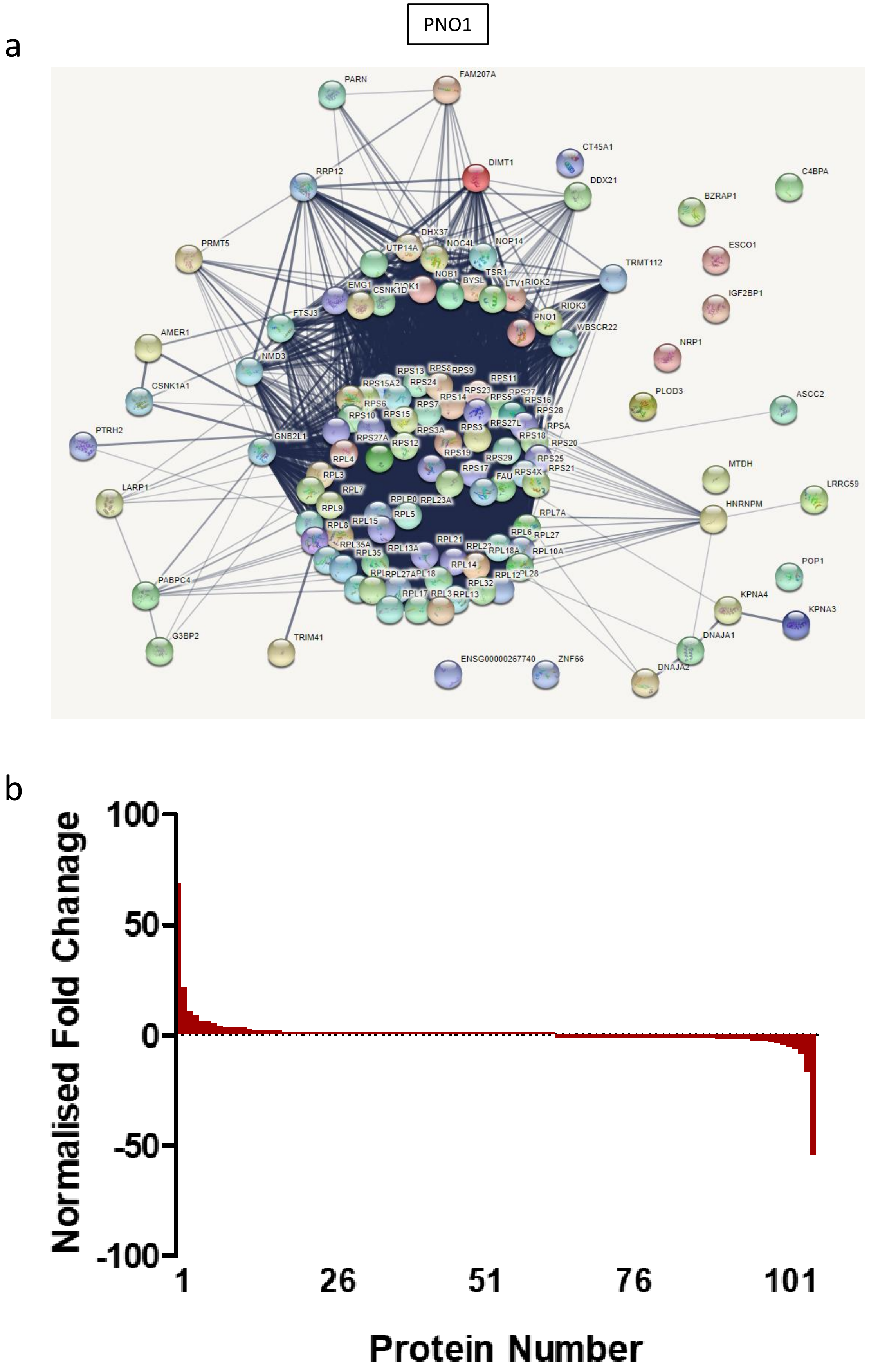


**Extended Data Figure 4. TMT LC-MS/MS analysis of PNO1 Twin-Strep-tag® pulldowns from a latent and 24 hour post lytic reactivation TREx BCBL1-Rta cell line.** STRING protein interaction map of identified proteins **(a)**. Fold change of all interacting proteins from latent compared to 24 hours post lytic reactivation **(b)**.

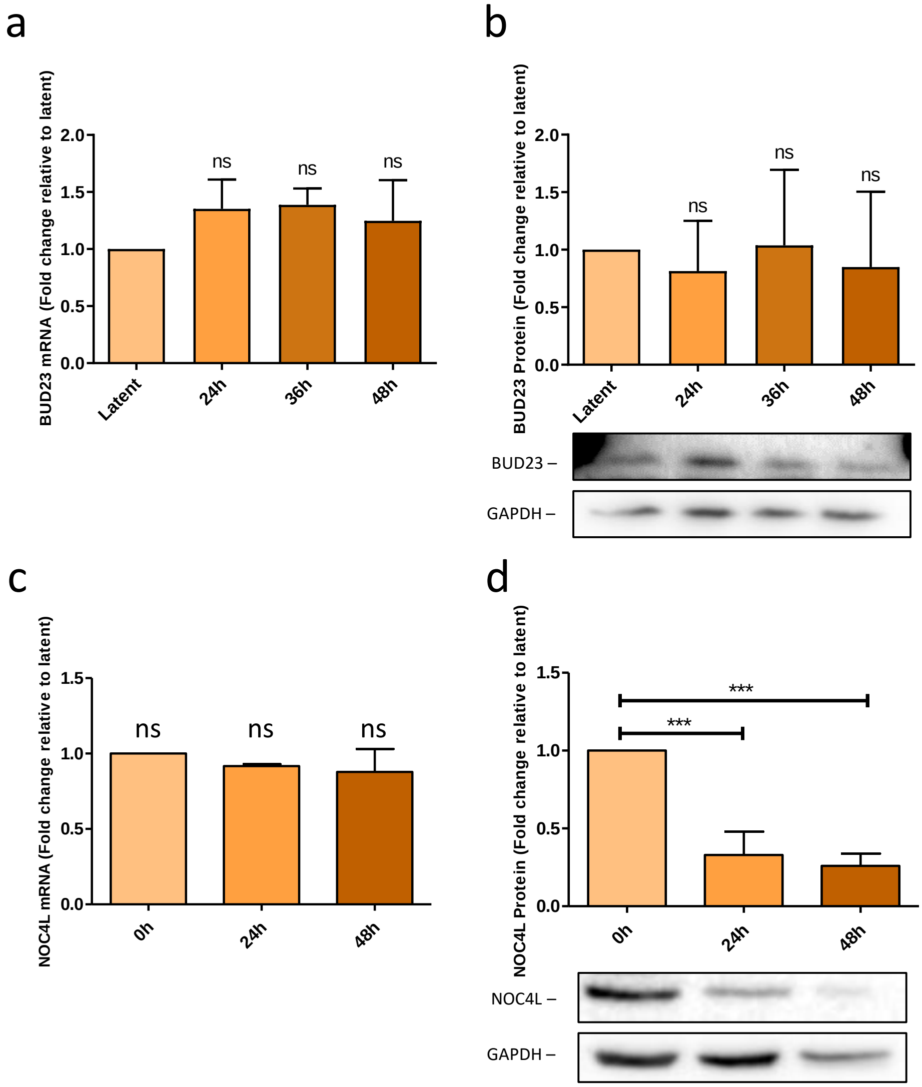


**Extended Data Figure 5. BUD23 and NOC4L expression during lytic reactivation of KSHV in TREx BCBL1-Rta cells.** Whole cell lysates were collected at various timepoints after lytic reactivation and split in two. Total RNA was isolated from half the cell lysates and BUD23 or NOC4L mRNA production was assayed by two step RT-qPCR and analysed by comparison to the 0h (latent) control using a ΔΔCt method (n=3-4) **(a,c)**. The second half of the whole cell lysates were analysed by western blot probing for BUD23 or NOC4L, GAPDH was included as a reference gene, representative western blots and densitometric analysis relative to the Scr control (n=3-4) **(b,d)**. Data are presented as mean ± SD. Significance was calculated by one-way ANOVA with a Newman-Keuls multiple comparison post-test. Asterisks denote a significant difference between the specified groups (* $p \leq 0.05$, ** $p<0.01$ and *** $p<0.001$). NS = Not significant.

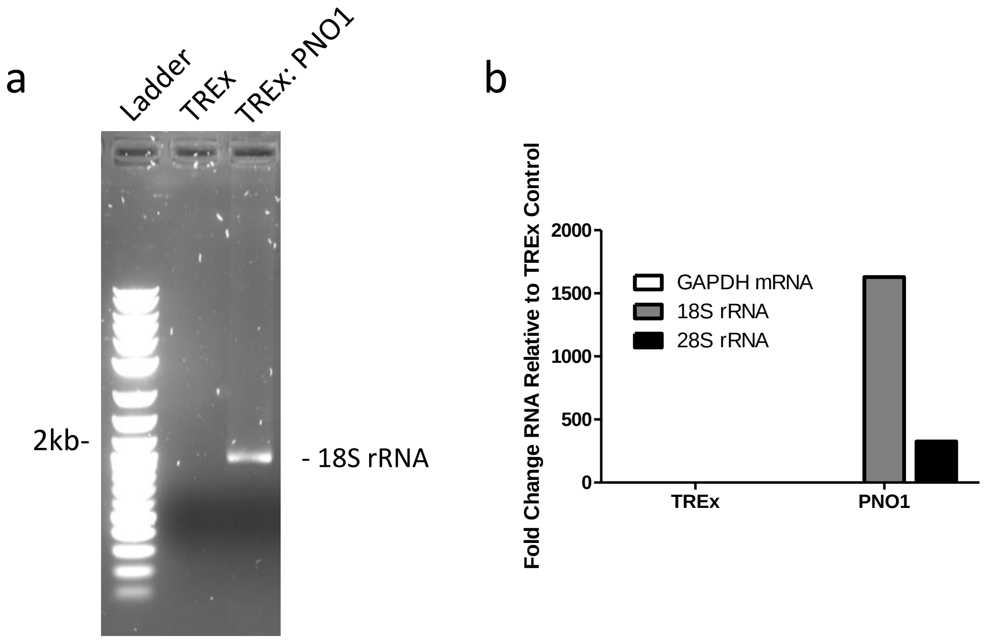


**Extended Data Figure 6. The RNA composition of isolated pre-40S complexes by PNO1 Twin-Strep-tag® pulldowns.** Total nucleic acids was isolated from whole cell lysate pulldowns from control cells and PNO1 bait protein expressing cells. Denaturing polyacrylamide gel electrophoresis **(a)** and two-step RT-qPCR, with primers specific for GAPDH mRNA, 18S rRNA and 28S rRNA **(b)**.

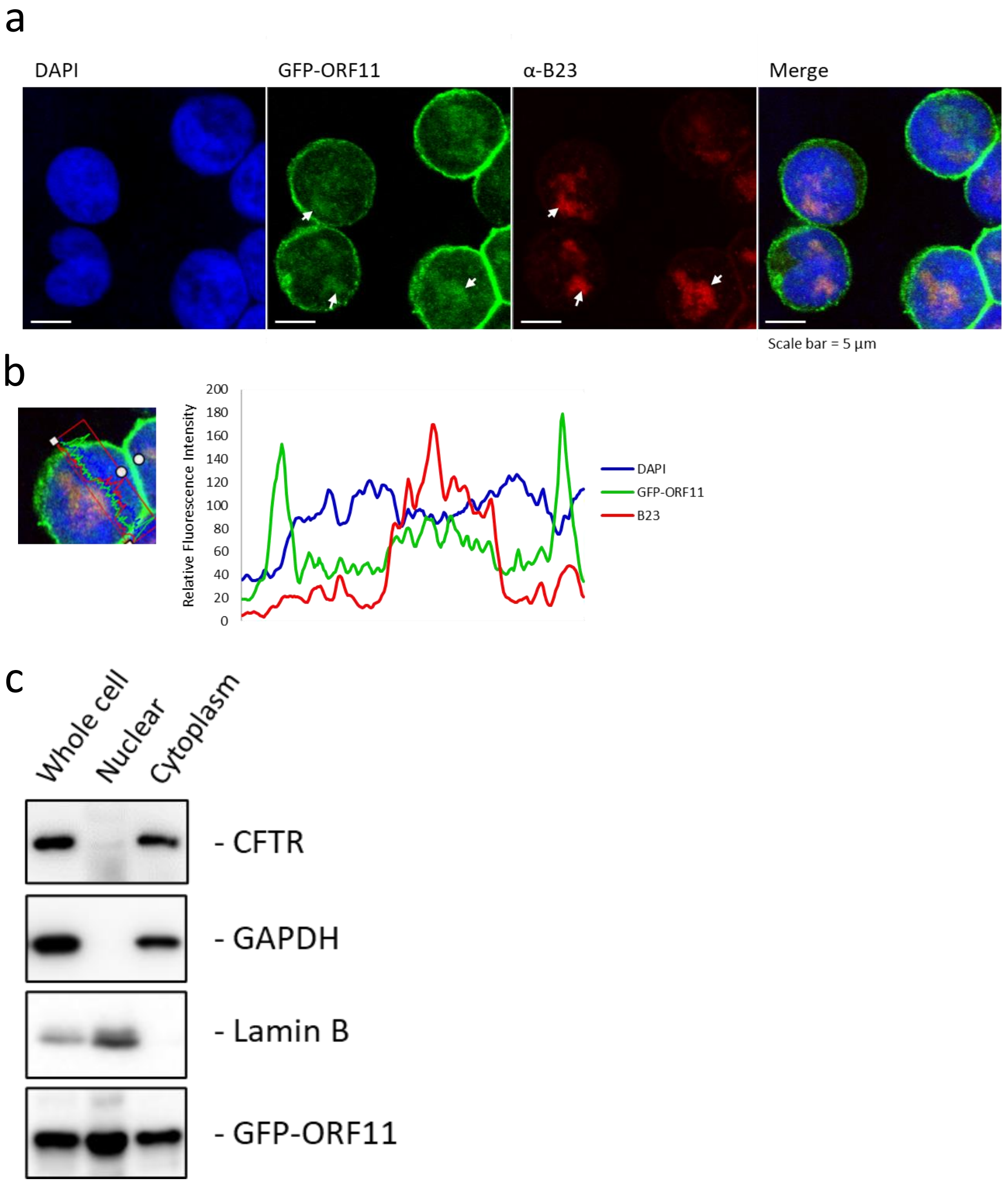


**Extended Data Figure 7. The KSHV protein ORF11 localises to the nucleolus and cell membrane.** KSHV lytic reactivation was induced for 8 hours in TREx BCBL1-Rta cells stably expressing GFP-ORF11 and were fixed, permeabilized, and stained for B23 (red) and the DNA dye DAPI (blue). The cells were mounted and viewed using an LSM 880 inverted confocal microscope **(a)**. A cross section of fluorescence intensity was calculated using ZEN blue 2.0 **(b)**. Lysates from TREx BCBL1-Rta cells stably expressing GFP-ORF11 were separated into nuclear and cytoplasmic fractions and analysed by western blot. Representative western blots are shown probing for a cytoplasmic/cell membrane marker CFTR, cytoplasmic marker GAPDH, nuclear marker Lamin B1, and GFP for GFP-ORF11 **(c)**.

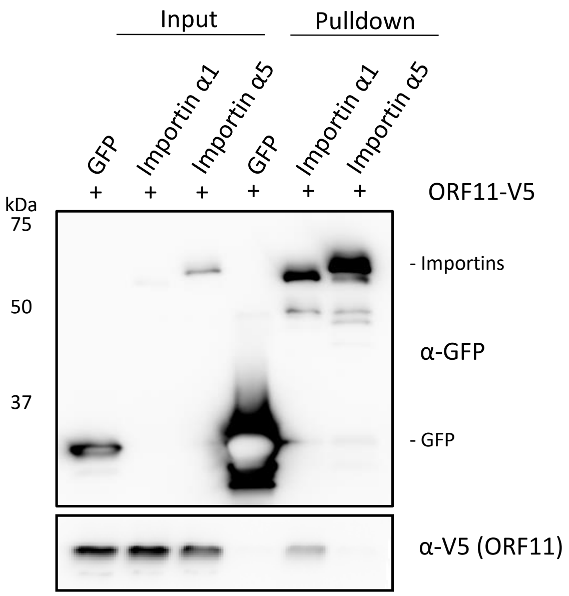


**Extended Data Figure 8. The KSHV protein ORF11 interacts with importin α1.** GFP-Trap® pulldowns of HEK 293T whole cell lysates. All cells were transfected with a plasmid encoding for the expression of ORF11-V5 and another plasmid encoding either GFP, GFP-importin α1 or GFP-importin α5.

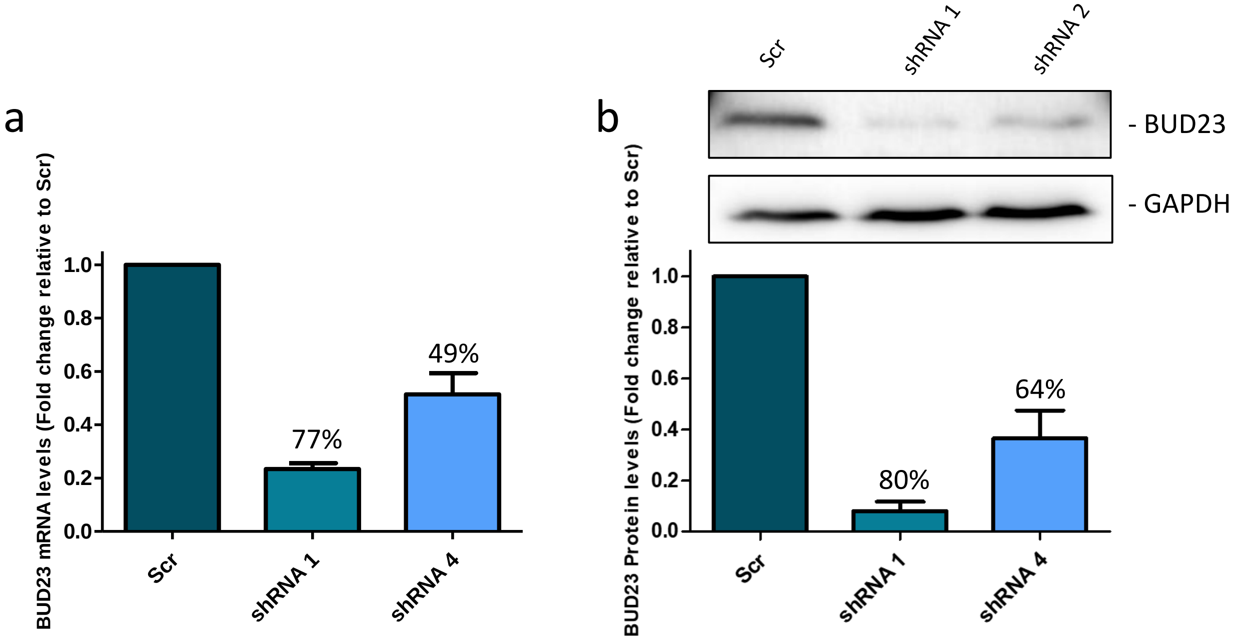


**Extended Data Figure 9. Knockdown of BUD23 in HEK 293T cells.** A Lentivirus expression system was used to stably transduced HEK 293T cells with a non-targeting scrambled shRNA (Scr) or two different shRNAs targeting BUD23 (shRNA 1 and 2). BUD23 mRNA production was assayed by two step RT-qPCR and analysed by comparison to the Scr control using a ΔΔCt method (n=4) **(a)**. Whole cell lysates were collected and analysed by western blot probing for BUD23, GAPDH was included as a reference gene, representative western blots and densitometric analysis relative to the Scr control (n=3) **(b)**. Data are presented as mean ± SD.

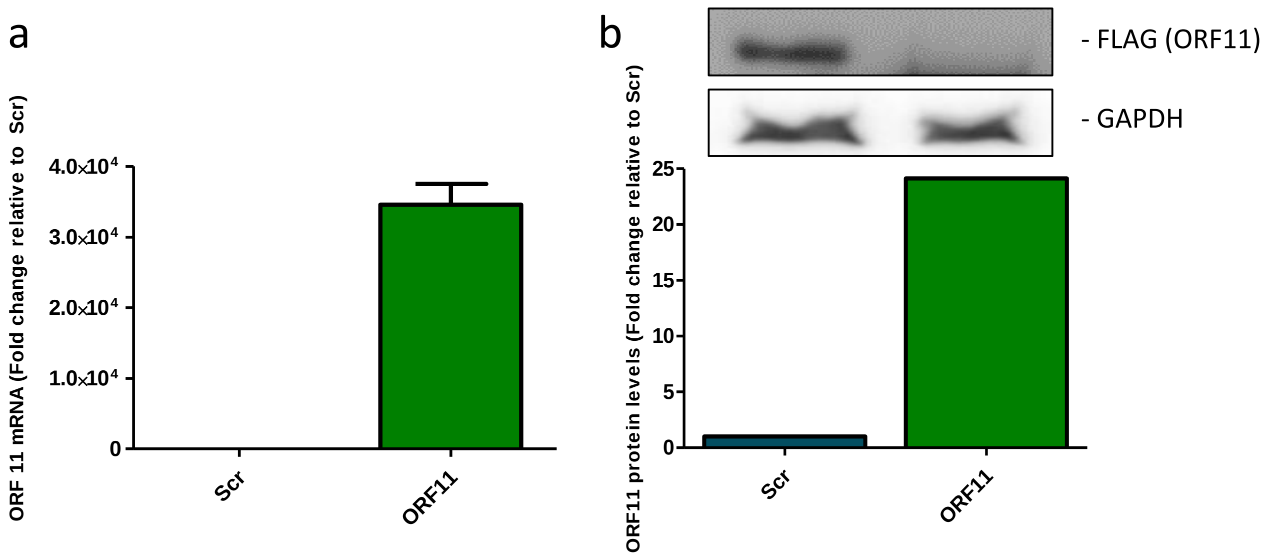


**Extended Data Figure 10. Expression of ORF11-FLAG in HEK 293T cells.** A Lentivirus expression system was used to stably transduced HEK 293T cells with a non-targeting scrambled shRNA (Scr) orORF11-FLAG expression plasmid. ORF11 mRNA production was assayed by two step RT-qPCR and analysed by comparison to the Scr control using a ΔΔCt method (N=2) **(a)**. Whole cell lysates were collected and analysed by western blot probing for FLAG, GAPDH was included as a reference gene, representative western blots and densitometric analysis relative to the Scr control (N=1) **(b)**. Data are presented as mean ± SD.

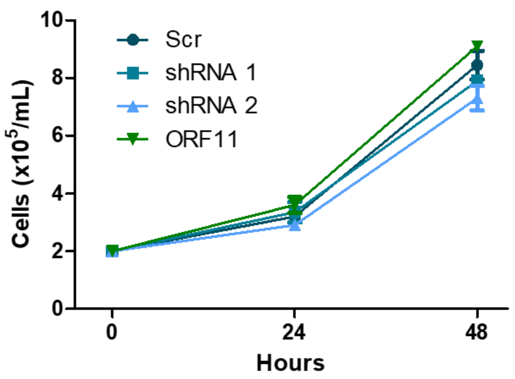


**Extended Data Figure 11. Proliferation of HEK 293T cell lines.** HEK 293T cell lines were counted over 48 hours to measure cell proliferation (n=2) Data are presented as mean ± SD.

a

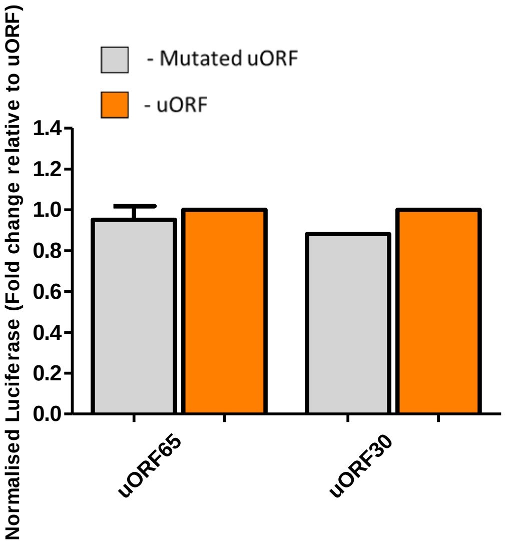


**Extended Data Figure 12. Luciferase reporter assay of KSHV uORFs which do not effect the expression of the downstream luciferase.** Normalised luciferase intensity was quantified from luciferase reported plasmids containing uORFs or with the start codon of the uORF mutated in HEK 293T cells (n=1-3). Data are presented as mean ± SD.

| Plasmid | Vector | Source |
|---|---|---|
| Lentiviral packaging | psPAX2 | A gift from Dr. Edwin Chen (University of Leeds) |
| Lentiviral envelope | pVSV.G | |
| FLAG-2xStep-DIMT1 | pCDH-CMV-MCS-EF1-Puro | GenScript (Custom made) |
| FLAG-2xStep-PNO1 | | |
| FLAG-2xStep-LTV1 | | |
| FLAG-2xStep-TSR1 | | |
| Scr shRNA | pLKO.1 | Dharmacon (RHS4459) |
| BUD23 shRNA 1 | | Dharmacon (TRCN0000140056) |
| BUD23 shRNA 2 | | Dharmacon (TRCN0000142391) |
| GFP | pLenti-III-mir-GFP | Cloned by aurthors for this study. |
| GFP-ORF11 | | |
| ORF11-FLAG | pLenti CMV GFP Puro | |
| ORF11-V5 | pcDNA™3.1/nV5-DEST | A gift from Prof. Yan Yuan (University of Pennsylvania) |
| GFP-importin α1 | pcDNAGFP | Previously described[47]. |
| GFP-importin α5 | | |
| Scr gRNA | lentiCRISPR V2 | Cloned by aurthors for this study. |
| ORF11 gRNA 1 | | |
| ORF11 gRNA 2 | | |

**Extended Data Table 1. All plasmids used in this study.**

| Primer | Forward | Reverse |
|---|---|---|
| GAPDH | TGT GGT CAT GAG TCC TTC CAC GAT | AGG GTC ATC ATC TCT GCC CCC TC |
| BUD23 | TAC GTT CGC AAC TCA CGG AT | CCA GCA GGT AAC AGG GCT TA |
| 18S rRNA total | GAT GGT AGT CGC CGT GCC | GCC TGC TGC CTT CTT TGG |
| 18S rRNA $m^7$G 1639 | GTA ACC CGT TGA ACC CCA TT | CCA TCC AAT CGG TAG TAG CG |
| ORF57 | GCC ATA ATC AAG CGT ACT GG | GCA GAC AAA TAT TGC GGT GT |
| ORF59 | CCG ATC GRG GAA AGG TAG GA | ATG TAC TCG ACG CTG GCA TA |
| K8.1 | GTTCCACACAGATTCGCACA | AGTTCATCCTGCCTAGCCAG |
| ORF65 | AAG GTG AGA GAC CCC GTG AT | TCC AGG GTA TTC ATG CGA GC |
| NOC4L | GCT TCT ATG TGA AGC GGG CG | GCC TGG AAA ACC CTC CTG TG |
| 28S rRNA | GGG TGG TAA ACT CCA TCT AAG G | GCC CTC TTG AAC TCT CTC TTC |
| ORF11 | TCC TCG AGC GTG CTG ATT TT | TAT TTT GAG CCC TCC CAC GG |

**Extended Data Table 2. All qPCR primers used in this study.** Displayed as 5'-3'.

| Target | Origin | Working Dilution | Supplier | Catologue # |
|---|---|---|---|---|
| GAPDH | Mouse | 1:5000 | Proteintech Europe | 60004-1-Ig |
| FLAG | Rabbit | 1:5000 | Sigma-Aldrich | 77425 |
| BUD23 | Rabbit | 1:500 | Thermo Fisher Scientific | PA521698 |
| NOC4L | Rabbit | 1:500 | Proteintech Europe | 17025-1-AP |
| RPS19 | Rabbit | 1:500 | Proteintech Europe | 15085-1-AP |
| ORF57 | Mouse | 1:1000 | Santa Cruz | sc-135747 |
| CDK1 | Mouse | 1:5000 | Abcam | ab18 |
| ORF59 | Rabbit | 1:1000 | A gift from Prof. Britt Glaunsinger (University of California, Berkeley) | N/A |
| K8.1 | Mouse | 1:1000 | Advanced Biotechnologies | 13-212-100 |
| ORF65 | Rabbit | 1:500 | Cambridge Research Biochemicals | crb2005224 |
| GFP | Mouse | 1:5000 | Proteintech Europe | 66002-1-Ig |
| DIMT1 | Rabbit | 1:500 | Proteintech Europe | 15563-1-AP |
| RPS3 | Rabbit | 1:500 | Proteintech Europe | 11990-1-AP |
| B23 | Mouse | 1:100 (IF) | Santa Cruz | sc-271737 |
| CFTR | Mouse | 1:500 | Santa Cruz | sc-376683 |
| Lamin B | Rabbit | 1:1000 | Abcam | ab16048 |
| V5 | Mouse | 1:1000 | Abcam | ab27671 |

**Extended Data Table 3. All antibodies used in this study for western blotting and immunofluorescence (IF).**
